# dmrff: identifying differentially methylated regions efficiently with power and control

**DOI:** 10.1101/508556

**Authors:** Matthew Suderman, James R Staley, Robert French, Ryan Arathimos, Andrew Simpkin, Kate Tilling

## Abstract

A differentially methylated region (DMR) is a genomic region in which DNA methylation is consistently positively or negatively associated with a phenotype or exposure. We demonstrate that existing algorithms for identifying DMRs either fail to consistently control false positive rates (comb-p and DMRcate), suffer from low power (bumphunter) or lack modeling flexibility (seqlm). We introduce a new method, dmrff, that overcomes these shortcomings and can additionally be used to meta-analyze multiple datasets. When applied to investigate associations of age in multiple datasets, dmrff identifies novel DMRs near genes previously linked to age. An R implementation is available on Github (http://github.com/perishky/dmrff).

## Background

DNA methylation is a modification of DNA by the addition of a methyl group. In mammals, methylation mainly occurs at cytosines, often in the context of a cytosine followed by a guanine (CpG). The presence of methylation is known to change how the underlying DNA sequence is interpreted within a cell (1). For example, methylation near the beginning of a gene is usually linked with lower gene activity (2). Patterns of DNA methylation are generally very stable and are copied faithfully from parent to daughter cell but do change throughout development and aging as well as with a variety of changes in phenotype and environmental exposures (3).

Epigenome-wide association studies (EWAS) test associations of DNA methylation levels at cytosines across the genome with phenotypes and exposures of interest. Most current EWAS are applied to DNA methylation datasets generated using either the Illumina Infinium HumanMethylation450 (450k) or MethylationEPIC (EPIC) BeadChips microarrays including measurements at 485,000 or 850,000 CpG sites, respectively. Although these are large numbers, they are a small fraction of the approximately 30 million CpG sites scattered across the human genome. To provide an informative coverage of the genome, measured CpG sites for both arrays were selected strategically to cover specific regions of interest including gene promoters, gene enhancers, CpG islands, transcription factor binding sites, and regions previously linked to disease. Each targeted region typically includes a small number of measured CpG sites. For example, over 15,000 genes have at least 5 measured CpG sites on the 450k array in their promoters. For the EPIC array, there are over 21,000 such genes.

Often EWAS studies lack power due to the curse of dimensionality, i.e. small numbers of samples relative to the number of measurements. To adjust for so many tests, most EWAS apply a simple Bonferroni-corrected threshold, 0.05 divided by the number of CpG sites tested. This threshold, however, is almost certainly conservative because it assumes independence between association tests at all pairs of CpG sites. However, there are strong dependencies between CpG sites, particularly CpG sites located close together. Using permutations, the recommended threshold was recently increased to 2.4×10^−7^ for the 450k array and 3.6×10^−8^ for genome-wide measurements (4).

A single threshold, however, cannot simultaneously account for the strong dependencies between specific groups of CpG sites, and lack of dependencies between others. A more sensitive approach for improving power may be to test associations with groups of dependent CpG sites, for example CpG sites located close together. In fact, associations are often observed at clusters of CpG sites within genomic regions (5) and are consequently called differentially methylated regions (DMRs). This is consistent with the role of DNA methylation making a genomic region either more or less amenable to binding by DNA binding proteins such transcription factors (6).

A variety of algorithms have been developed for detecting DMRs including bumphunter (5), Comb-p (7), DMRcate (8) and seqlm (9). Each method other than seqlm begins with an EWAS and then adjusts CpG site summary statistics by sharing information between nearby CpG sites. Candidate DMRs are identified as regions composed of CpG sites whose adjusted statistics all surpass some threshold. Finally, statistics are calculated for each candidate DMR. Seqlm differs in that it first partitions the genome into regions whose CpG sites have similar associations with the variable of interest. DNA methylation summaries of each region are then generated and an EWAS is applied to these region summaries.

Other DMR-finding algorithms have been proposed but have been omitted from our comparison for various reasons. Aclust (10) is omitted because it is known to generate a high number of false positives (9). Several other methods are omitted because they only support comparison between two groups: GetisDMR (11), DMRFinder (12), Probe Lasso (13) and DMRMark (14). Recently proposed GlobalP (15) and SKAT (16) were found to by their authors to produce inflated statistics in simulated data. Originally designed for genome-wide association studies, aSPUw was recently applied to DNA methylation data (17). We omit aSPUw from our comparison because it considers a different DMR model, e.g. allowing direction of effect to differ between CpG sites.

In the following, we investigate the performances of bumphunter, Comb-p, DMRcate and seqlm and identify important shortcomings of each. We propose a new method, dmrff, that overcomes each of these shortcomings.

## Results

Figure 1 illustrates DMRs identified by dmrff in the meta-analysis 14 publicly available datasets to identify associations with age (see Methods). All DMRs lie within the *BLCAP* gene, near the beginning of the *NNAT* gene. The extent of each is marked by a dark gray horizontal bar. Meta-analysed effect estimates are shown as thin vertical bars for each measured CpG site. Two DMRs are composed of CpG sites with negative effects and a third much larger DMR is composed of CpG sites with positive effects.

**Figure 1.**
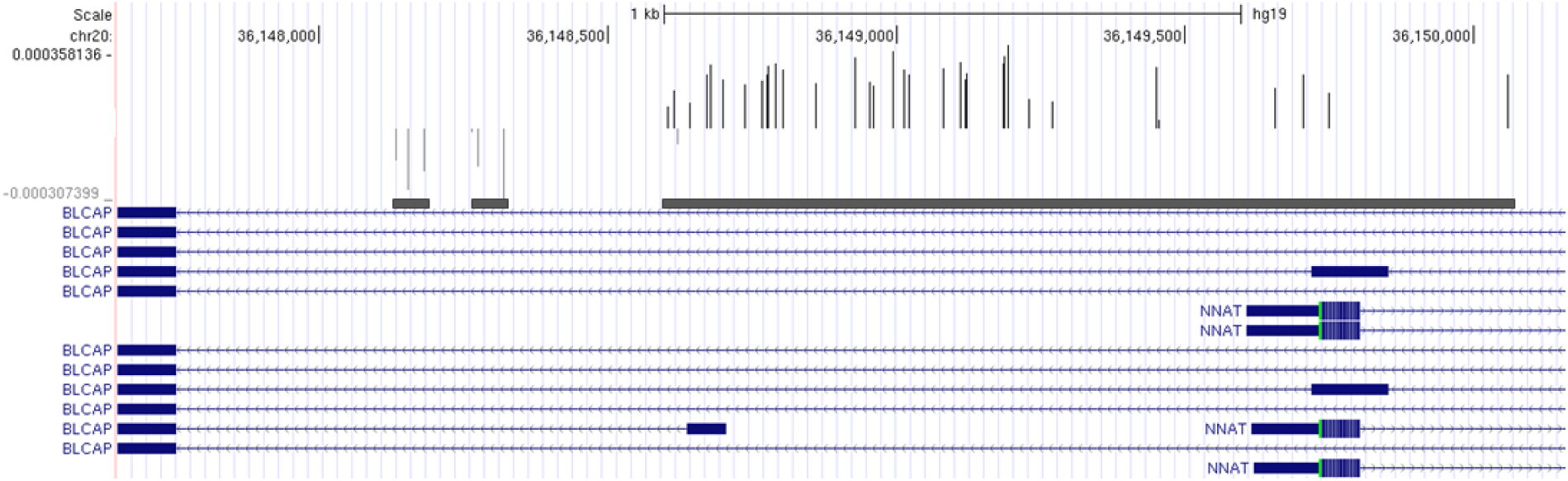
Plot of differentially methylated regions (DMRs) associated with age. Plot shows EWAS t-statistics for CpG sites as a bar graph, one bar for each CpG site (dark gray bars show positive statistics, light gray bars show negative) in the UCSC Genome Browser (25). Differentially methylated regions are identified by thick, dark gray horizontal bars. Transcripts of the BLCAP gene are shown in blue.

### Simulation Studies

We applied algorithms comb-p, DMRcate, bumphunter and seqlm as well our new algorithm, dmrff, to two different simulations: one for estimating false positive rates and another for estimating power (see Methods).

### False positive rates

False positive rates (FPRs) were estimated by generating random phenotype/exposure variables and testing associations with publicly available DNA methylation profiles (see Methods for details). All maintained false positive rates below expected 5% threshold except for comb-p (FPR > 20%; Figure 2) when applied to a subset of chromosome 1 (‘chr1-subset’, see Methods). However, when applied to a subset of the genome including clusters of highly correlated CpG sites (‘corr-subset’, see Methods), we found that the bumphunter ‘value’ statistic was also slightly inflated (FPR = 8%; Figure 2). We also found that DMRcate statistics were prone to inflation but was not observed unless the dataset contained at least one CpG site association with a false positive rate below 0.05. When we tweaked the dataset by adding additional CpG sites to guarantee at least one such association (‘augmented corr-subset’, see Methods), we found that the false positive rates for both comb-p and DMRcate increased to 91.7% and 51.6%, respectively (Figure 2). Note that any DMRs identified that included the additional CpG were not included in the false positive rate calculations. The dramatic false positive increase for comb-p is due to the additional CpG sites having lower inter-CpG correlations than in the rest of the dataset. This caused the comb-p auto-correlation function to under-estimate correlations between CpG sites leading to greater inflation of DMR statistics. Only the bumphunter ‘area’ statistic, EWAS, seqlm and dmrff maintained false positive rates below the 0.05 significance threshold used by each algorithm.

**Figure 2.**
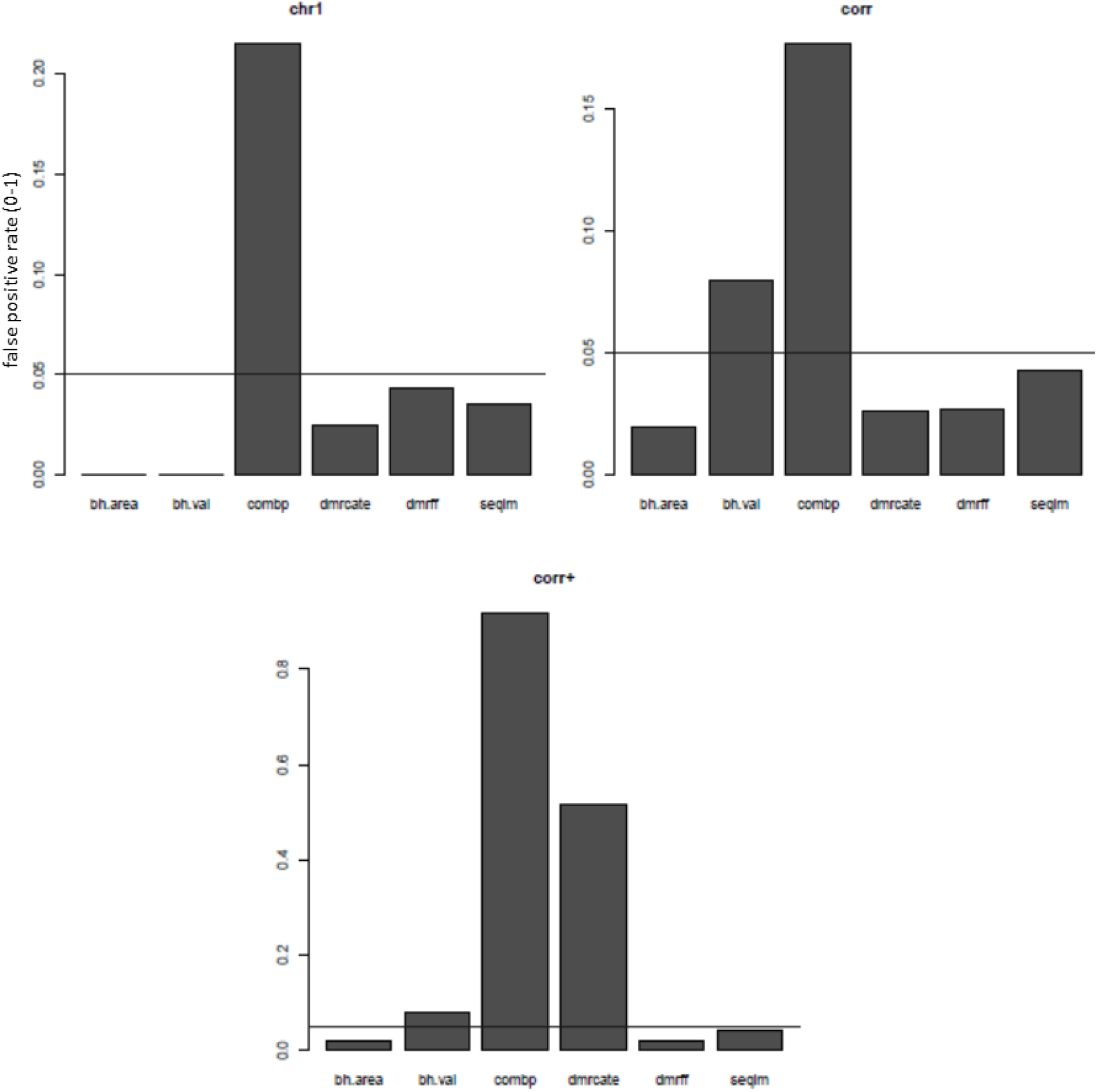
False positive rates of DMR algorithms. Rates are provided for three datasets, chr1, corr and corr+ (see Methods for details). The p-value threshold (adjusted for multiple tests) was set at 0.05 so any rate above 0.05 (horizontal line) indicates failure to control false positive rates.

### Power

The power of each DMR algorithm was assessed by generating phenotype/exposure variables associated with clusters of CpG sites in the methylation dataset at various strengths (see Methods). Power at each association strength was estimated as the proportion of generated variables for which an algorithm identified a DMR that overlapped with the associated cluster of CpG sites. Overall, comb-p has the greatest power followed by DMRcate, seqlm and dmrff, EWAS and finally the two versions of bumphunter (Supplementary Table 1). Figure 3 shows the power of only seqlm, dmrff and EWAS because power lacks meaning for algorithms that do not control false positive rates.

**Figure 3.**
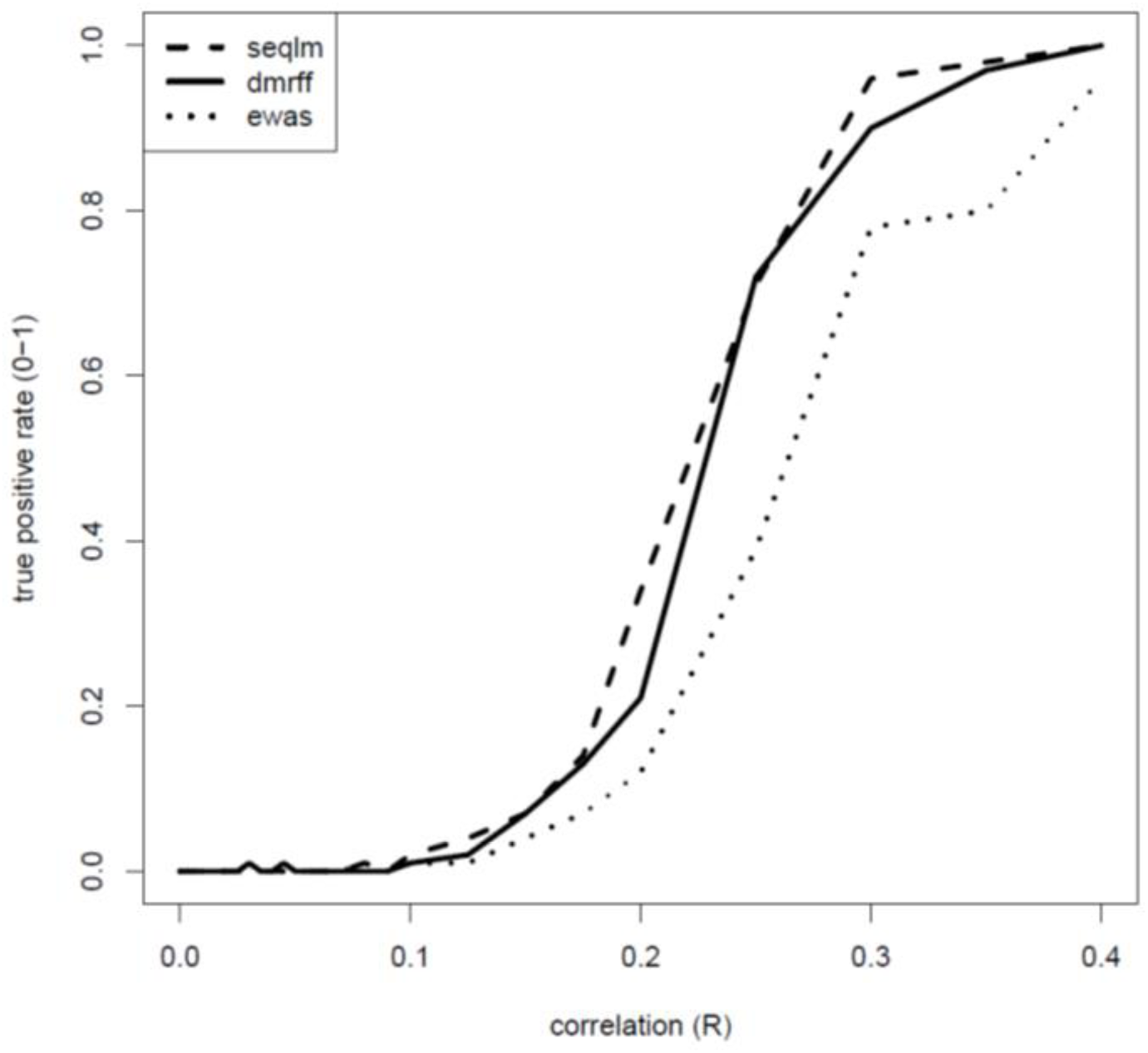
True positive rates of DMR algorithms. Shown is the proportion of simulated DMRs identified by the algorithms (y-axis) at the given strength of association between the CpG sites in the simulated DMRs and the variable of interest (x-axis). See methods.

### Analysis of age

We applied dmrff, seqlm and EWAS to an analysis of age in 14 publicly available DNA methylation datasets (see Methods). In each dataset, all three methods identified similar numbers of DMRs although numbers varied quite significantly between datasets (Supplementary Table 2). DMR numbers were estimated from EWAS by merging CpG sites with age associations within 500bp of one another into a single DMR. Overall, dmrff appears to have consistently identified more DMRs than the other two methods. To investigate if this is sensitive to significance thresholds, we asked how many DMRs remained for each method after removing any that overlapped with DMRs identified by another method but at a more relaxed significance threshold (Supplementary Table 3). Both dmrff and seqlm identified hundreds of DMRs (at Bonferroni-adjusted p < 0.05) containing no EWAS associated CpG sites (at Bonferroni-adjusted p < 0.2). dmrff identified hundreds and, for over 70% of the datasets, thousands of DMRs (at Bonferroni-adjusted p < 0.05) that did not overlap with DMRs identified by seqlm (at Bonferroni-adjusted p < 0.2). seqlm identified at most 326 DMRs (at Bonferroni-adjusted p < 0.05) in any single dataset not overlapping with any identified by dmrff (at Bonferroni-adjusted p < 0.2).

### Replication of DMRs

We asked how well DMRs identified by dmrff in one dataset replicated in other datasets. For each of the 14 publicly available datasets described above, we meta-analysed the statistics for the age DMRs identified by dmrff in the remaining 13 datasets (fixed-effect inverse-variance weighted meta-analysis) and then calculated the rate of replication (i.e. the % of DMRs with meta-analysed Bonferroni adjusted p < 0.05). For comparison, we used the same approach to calculate replication rates for individual CpG sites. The results are summarized in Supplementary Table 4. In 12 out of 14 datasets, we observed above 96% replication rates and less than 0.13% difference between the CpG site and the DMR replication rates in each dataset. The two remaining dataset replication rates were 1.2% and 3% higher for CpG sites.

We also calculated replication rates for DMRs that did not contain age-associated CpG sites (at Bonferroni-adjusted p < 0.2). For 11 of 14 datasets, the CpG site replication rate was less than 2% greater. For the remaining three datasets, the CpG site replication was 20%-37% greater. For two of these datasets, GSE55763 and GSE87571, CpG site replication rates, 87.4% and 61.2%, respectively, was unusually low compared to other datasets. In all other datasets, replication rates were above 96%. This suggests that these two datasets are somehow systematically different from the other datasets.

### Meta-analysis

It is now common to increase statistical power by meta-analyzing summary EWAS statistics from multiple studies. dmrff can be extended for use in meta-analysis in a straight-forward way using two different approaches, the first called ‘two-step meta-analysis’ and the more convenient ‘reference meta-analysis’ (see Methods). In both approaches, EWAS summary statistics are calculated for each dataset, meta-analysed and used to identify candidate DMRs. Candidate DMR statistics are then calculated for each dataset using EWAS summary statistics and finally meta-analysed across all datasets. In the two-step approach, pairwise CpG site correlations used to calculate candidate DMR statistics are obtained from each original dataset. In the reference approach, the pairwise correlations are obtained from a single designated dataset.

In the 14 datasets described above, the two-step approach identified 70,464 DMRs of age whereas the reference approach identified 83,549. However, this difference however is misleading since the two step approach generally identifies much larger DMRs (Supplementary Table 5). For example, 22,973 two-step DMRs contain more than 1 CpG site compared to only 12,156 reference DMRs. In addition, the two-step DMRs are much more likely to be novel (Supplementary Table 5). A DMR is considered novel for an approach if it contains no CpG sites belonging to a DMR identified by the other method (at Bonferroni adjusted p < 0.2 rather than 0.05). For example, of the 9,095 two-step DMRs covering at least 8 CpG sites, 347 are novel. By comparison, there are only 204 DMRs covering 8 or more CpG sites, of which only 9 are novel. For most DMR size thresholds, the two-step approach identifies at least 10x more novel DMRs than the reference procedure. Since CpG correlations play a larger role in the DMR statistic for larger DMRs, these results indicate that a reference dataset is unlikely to provide a useful representation of the CpG site correlation structure found in each individual dataset.

### Novel DMRs are functionally relevant

Of the 70,464 age DMRs identified by the two-step method, 662 do not include any CpG site association identified by EWAS meta-analysis (Bonferroni-adjusted p < 0.05). Relaxing the EWAS p-value threshold from 0.05 to 0.2, 482 remain without an EWAS association.

We investigated the potential biological implications of these 482 novel DMRs by comparison to published associations of gene expression profiles from nearly two hundred individuals in nine tissues: adipose, artery, heart, lung, muscle, nerve, skin, thyroid and blood (18). Of 41298 genes included in the gene expression profiles, 14308 were linked (by Illumina annotation) to one of our age DMRs. Eliminating all those genes that were also linked to an age-associated CpG site identified by EWAS meta-analysis (at a relaxed Bonferroni-adjusted p < 0.2), 123 DMR-linked genes remained.

We then asked if these 123 genes were enriched with gene expression associations with age. We observed strong enrichments in blood but not in any other tissues. For example, associations with age had been observed in 51 (40%) of these 123 genes in blood but at most 21 (17%) in any of the other eight tissues (adipose 14, artery 21, heart 6, lung 4, muscle 5, nerve 0, skin 0, thyroid 0).

To formally assess enrichment, we applied two enrichment tests. First, we used Fisher’s exact test with respect to several different significance thresholds for identifying genes with age-associated gene expression levels (p = 1 × 10^−6^-0.2). We observed very strong enrichments in blood for most thresholds considered, but no evidence of enrichment in any of the other tissues (Figure 4). Second, we applied the Wilcoxon rank sum test to determine if the 123 genes were more strongly associated with age than a random selection of the same number of genes. Again, in blood, we observed strong evidence of enrichment (p < 0.0012) but no evidence in the other tissues (p > 0.26).

**Figure 4.**
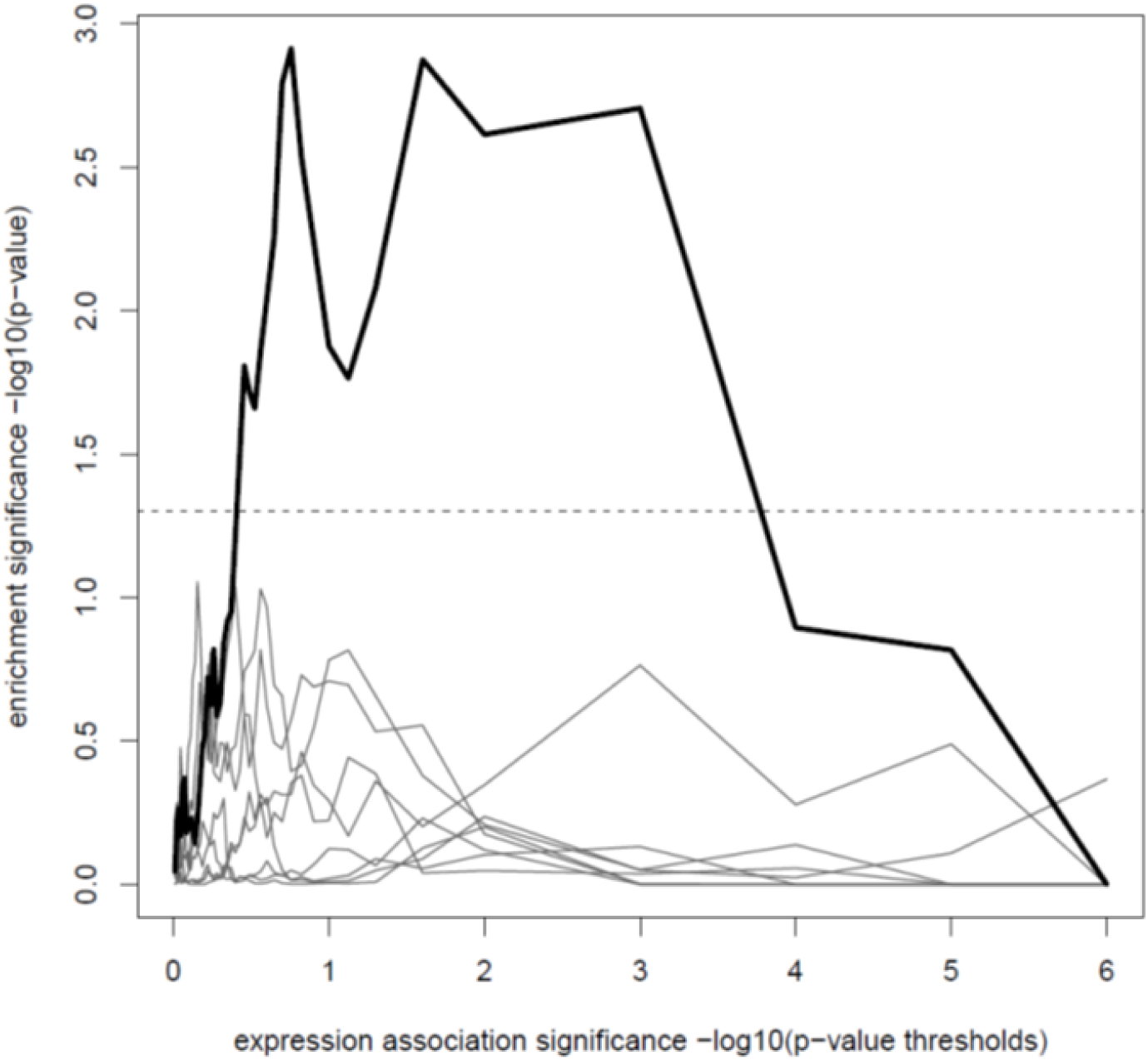
Enrichment of gene expression associations with age among genes linked with novel age DMRs. The 123 genes linked to DMRs of age but not to CpG sites associated individually with age were tested using Fisher’s exact test for enrichment with genes associated with age in nine tissues: adipose, artery, heart, lung, muscle, nerve, skin, thyroid and blood. Shown is the statistical significance of Fisher’s exact test (y-axis) for different p-value thresholds for identifying gene expression associations with age.

### Running time

Comb-p, DMRcate, seqlm and dmrff are all complete computation of a genome-wide dataset in a few minutes, comparable to the running time of an EWAS (Table 1). Bumphunter requires about 10x longer when using 100 bootstraps.

**Table 1.**
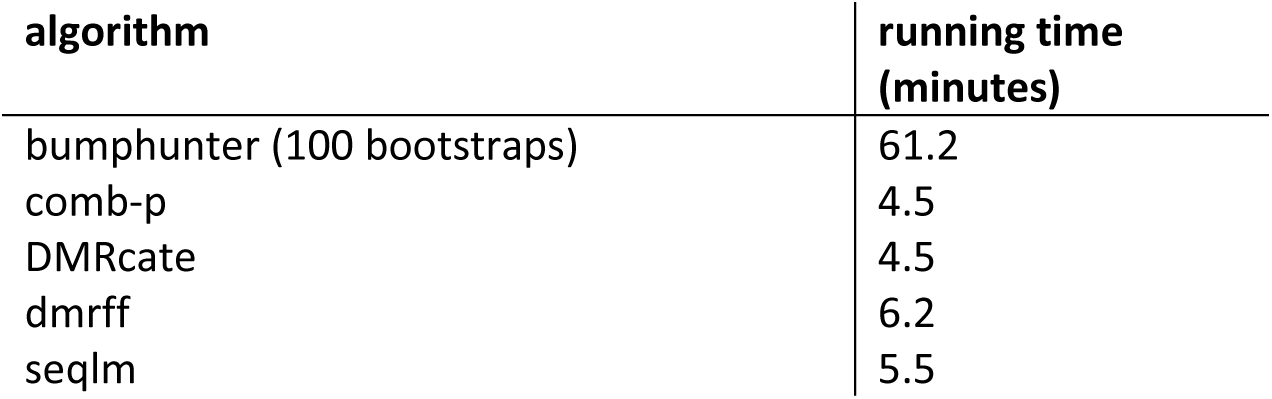
Running times of algorithms applied to DNA methylation profiles including 470K CpG sites and 464 samples with 44 covariates on a computer with 10 processors.

## Discussion

We have described a new approach for identifying differentially methylated regions based on inverse-variance weighted meta-analysis that is fast, controls false positive rates and is often more powerful than other approaches like EWAS and seqlm that also control false positive rates. Furthermore, when applied to several publicly available datasets, we show that DMRs identified are highly replicable between datasets and that novel DMRs identified by our extension of dmrff for meta-analysis are biologically relevant.

None of the previously published DMR-finding algorithms can make all these claims. Comb-p is one of the most popular algorithms because it is extremely powerful and was one of the first serious attempts to avoid false positives due to dependencies between CpG sites. Unfortunately, we and others (9) have shown that the approach fails to consistently control error rates because it depends on a single autocorrelation function to represent a DNA methylation correlation structure that varies across the genome. Consequently, in some genomic regions, the function underestimates dependencies between CpG sites and consequently generates inflated DMR statistics.

DMRcate is also very popular due to its speed and ease of use. It is, however, often not more powerful than EWAS because it tests for DMRs only when EWAS successfully identifies associations. Furthermore, similar to comb-p, it tends to generate inflate DMR statistics in genomic regions with strong dependencies between CpG sites.

Although bumphunter is an intuitive approach for identifying differentially methylated regions and one of the earliest approaches to allow the data to completely determine the beginning and end of DMRs, it is unfortunately extremely time-consuming and lacks power. Further, we have shown that the ‘value’ statistic may also be inflated in parts of the genome with strong dependencies between CpG sites.

A recently published method, seqlm, is currently the only that correctly controls false positive rates and may additionally be more powerful than EWAS. However, we show many examples where it is less powerful than dmrff. More importantly, its implementation is not as flexible as that of dmrff, as it is limited to a single association-testing model and does not appear to be have a straight-forward adaptation for meta-analysis.

Dmrff does have some limitations. First, it does sacrifice a small amount of power in order to optimize the starting and ending positions of DMRs. Candidate regions are initially identified as sequences of CpG sites with EWAS p-values below some threshold, e.g. 0.05. Dmrff then shrinks the regions by calculating statistics for all sub-regions and then greedily selecting sub-regions to cover the candidate region with the strongest statistics. This sub-region analysis increases the burden of multiple testing.

Second, the adaptation of dmrff to meta-analysis requires CpG site correlations from each of the original datasets. In meta-analyses performed in large consortia, a straight-forward application of dmrff involves a somewhat clumsy two-step process in which dataset analysts first perform EWAS and then wait for EWAS meta-analysis and candidate DMR discovery to be performed centrally before going ahead with calculating statistics for candidate DMRs. To make meta-analysis more convenient, we have implemented a procedure that generates and saves the correlations of all pairs of CpG sites a given maximum distance *δ* apart in a dataset. This procedure is applied to each dataset at the same time as the original EWAS. These correlations can then be submitted at the same time as the EWAS summary statistics for meta-analysis. It is then possible to perform the remaining dmrff meta-analysis centrally without further consulting individual datasets. The DMRs identified using this approach are the same as for the basic two-step procedure provided that no candidate regions are larger than 2*δ* base pairs in length. However, in some meta-analyses, original datasets and therefore CpG site correlations are no longer available such as when previously published studies are meta-analysed. In these cases, dmrff can still be applied using correlations from a reference datataset. However, as we have shown, the result will be less powerful. We note that this is still preferable to using algorithms like comb-p or DMRcate that do not require CpG site correlations, because the results are likely to suffer from inflated p-values.

Third, dmrff is not currently the best option for whole-genome bisulfite sequencing data which includes DNA methylation measurements for all or nearly all the CpG sites in the genome. This is because the greedy approach for shrinking candidate genomic regions will likely generate far too many tests in genomic regions with high CpG density. In future, we plan to investigate more efficient replacements for the greedy algorithm.

## Conclusions

We recommend dmrff as the best of available options for identifying DMRs in DNA methylation profiles generated using the Illumina BeadChips. An open-source implementation is freely available on GitHub (http://github.com/perishky/dmrff).

## Methods

### Published algorithms for identifying DMRs

Comb-p (7), DMRcate (8), bumphunter (5) and seqlm (9) were all applied with default settings with the following exceptions. Comb-p was applied with dist set to 500, seed to 0.05 and threshold to 0.05. DMRcate was applied with lambda set to 500 and min.cpgs set to 2. Bumphunter was applied with useWeights and pickCutoff set to ‘TRUE’, pickCutoffQ to 0.95, nullMethod to ‘bootstrap’, B to 100 and maxGap to 500. Seqlm was applied with max_block_length set to 20 and max_dist to 500. To reduce running time of bumphunter, in cases where other algorithms were applied to 1000 simulations, bumphunter was applied to 100.

Each of these algorithms except for seqlm begins with an EWAS and then adjusts CpG site summary statistics by sharing information between nearby CpG sites (500bp here). Candidate DMRs are identified as regions composed of CpG sites whose adjusted statistics all surpass some threshold. Finally, statistics are calculated for each candidate DMR.

Bumphunter smooths CpG site EWAS effects using local regression with a Gaussian kernel (by default but other approaches can be specified). Candidate DMRs are identified as sets of consecutive CpG sites whose smoothed effects are all greater than a pre-specified threshold. Statistical significance is calculated by generating a null distribution by bootstrapping.

Comb-p operates on EWAS p-values, updating the p-value for each CpG site by combining p-values of nearby CpG site using the Stouffer–Liptak–Kechris (SLK) correction and then adjusting the resulting p-values for multiple tests. In the correction, p-values are weighted by their distance from the CpG site whose p-value is being updated according to an auto-correlation function derived from the EWAS p-values. Candidate DMRs are identified as sets of consecutive CpG sites with updated p-values below some threshold. P-values for candidate DMRs are then again calculated using the SLK correction and the autocorrelation function.

DMRcate smooths EWAS t-statistics by combining t-statistics from nearby CpG sites using a Gaussian kernel of a specified width. Statistical significance for each CpG site is then recalculated from the smoothed t-statistic and a new threshold for genome-wide significance selected so that the number of CpG sites passing the threshold is the same as the number that survived multiple testing correction in the original EWAS. Consecutive CpG sites whose recalculated p-values are below this threshold are identified as DMRs.

Seqlm differs from these algorithms by first partitioning the genome into regions of CpG sites with similar associations with the variable of interest. Region boundaries are identified using the minimum description length (MDL) principle. Following this, a DNA methylation summary is obtained for each region and then tested for associations with the variable of interest.

### dmrff

Similarly to bumphunter, comb-p and DMRcate, dmrff identifies DMRs by combining EWAS summary statistics from nearby CpG sites. Straight-forward approaches for combining summary statistics such as Fisher’s method or other meta-analytic approaches cannot be directly applied because they assume independence between underlying tests. In our case, this assumption corresponds to independence between nearby CpG sites which is rarely true. Violating this assumption causes combined statistics to have inflated statistical significance. An extension of inverse-variance weighted meta-analysis was recently proposed for such cases (19), and we apply it here to EWAS statistics for the CpG sites within a genomic region. The test statistic is B/S where

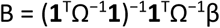

and

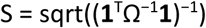

and β is the one-column matrix of EWAS effect estimates for each CpG site, Ω is equal to σσ^T^ x ρ, σ is the vector of standard errors for β, ρ is the CpG site correlation matrix, and **1** is a one-column matrix of 1’s. The test statistic B/S for the region with null associations has a standard normal distribution.

Genomic regions to test are identified as those spanned by sets of CpG sites with at most 500bp between consecutive sites and which have nominal EWAS p-values < 0.05 and EWAS effect estimates with the same sign (all of these values are parameters in dmrff and can be redefined). The region statistic B/S is then calculated not only for each region but also for each sub-region. Candidate DMRs are identified by greedily selecting sub-regions with the most extreme statistic that do not overlap with previously selected candidates. P-values for each candidate are Bonferroni-adjusted for multiple tests, conservatively treating each EWAS test and each sub-region test as independent.

Inputs to dmrff include the EWAS effect size estimates and p-values along with the EWAS DNA methylation profiles. For our analyses, limma (20) was used to execute the EWAS.

### Simulations

We generated simulations to evaluate the power and false positive rates of DMR algorithms. To ensure that the simulations were as similar to real-world applications, we produced simulations from DNA methylation profiles generated from human blood DNA using the Illumina Infinium HumanMethylation450 Beadchip. The 464 profiles are publicly available at the Gene Expression Omnibus (accession number GSE50660) (21). We applied algorithms to quite small subsets of the 458K CpG site measurements available from each profile in order to decrease algorithm running time per simulation. This allowed us to increase the precision of performance estimates by increasing the number observations per algorithm. We reduced the 485K CpG site measurements across the genome to three different subsets:

1. ‘chr1-subset’: The 2577 autosomal CpG sites found in clusters of 15-25 measured CpG sites on chromosome 1. A cluster was defined as a sequence of measured sites with less than 500bp between consecutive CpG sites. These data represent medium-sized clusters of sites.
2. ‘corr-subset’: The 8906 autosomal CpG sites found in clusters of at least 2 CpG sites in which consecutive measured CpG sites had a correlation of R > 0.8. These data represent (potentially) smaller sized groups of sites with high correlation between sites used to define clusters.
3. ‘augmented corr-subset’: The corr-subset of 8906 CpG sites on chromosome 1 together with CpG sites with simulated methylation levels correlated with the variable of interest. These additional CpG sites are needed to reveal inflated DMR statistics generated by DMRcate. When DMRcate is applied, if no individual CpG site has an association with the variable of interest with a false discovery rate less than 5%, then DMRcate terminates and reports no DMRs. However, if even a single CpG site has a sufficiently strong association, then DMRcate proceeds to test for DMRs and reports statistics for each candidate DMR. We therefore artificially included CpG sites with sufficiently strong associations with the variable of interest. To prevent these additional sites from interfering with DMRs corr-subset and to satisfy the DMRcate requirement that all CpG sites correspond to those measured by the Illumina microarray, we generated random methylation levels for the 670 CpG sites located in clusters of 10 CpG sites on chromosome X. Correlations of these CpG sites with the variable of interest were uniformly distributed between R=0 and R=0.3. DMRs identified by any algorithm on chromosome X were omitted from error rate calculations.

The full simulation dataset consisted of the selected DNA methylation dataset subset, a simulated exposure/phenotype variable and covariates obtained from the original dataset. Covariates included smoking, age and sex because they are known to have strong associations with many CpG sites throughout the genome.

The simulated exposure/phenotype variable was generated from the selected DNA methylation dataset subset. The variable was chosen to be continuous and drawn from the standard normal distribution (i.e. mean 0, standard deviation 1). Simulation details are described below.

#### Null simulations

Null simulations consisted of 1000 random variables drawn from a standard normal distribution. These were generated independently of the methylation data or covariates, so they could be used to estimate the false positive rate of an algorithm. Specifically, the false positive rate would be equal to the number of variables for which the algorithm identified at least one DMR.

#### Power simulations

A phenotype/exposure variable with a differentially methylated region in a methylation dataset was simulated by:

1. identifying a cluster of at least 15 CpG sites (at most 500bp between consecutive CpG sites),
2. taking the mean of the standardized methylation levels of the 10 CpG sites in the middle of the cluster, and then
3. generating a random variable with a given correlation R with the mean.

By construction, the resulting variable would have an average correlation of approximately R with each of the 10 consecutive CpG sites.

The correlations considered ranged from 0 to 0.4:

0.0, 0.005, 0.01, 0.015, 0.02, 0.025, 0.03, 0.035, 0.04, 0.045, 0.05, 0.06, 0.07, 0.08, 0.09, 0.1, 0.125, 0.15, 0.175, 0.2, 0.25, 0.3, 0.35, 0.4

For each of the 24 possible correlations, 100 variables were generated.

The power of an algorithm at a given correlation is then estimated as the proportion of the 100 simulated variables for which the algorithm identifies a DMR overlapping with at least one of the 10 CpG sites used to generate the variable.

### DNA methylation datasets for analyses of age

We performed analyses of age in publicly available DNA methylation profiles of peripheral blood in order to test the methods in a real-world setting. We identified 20 eligible datasets available from the Gene Expression Omnibus (GEO) (22), all generated using the Illumina Infinium HumanMethylation450 BeadChip (Supplementary Table 6). An EWAS of age was performed in each using linear models implemented in the limma R package (20). Chromosomes X and Y were excluded as datasets included males and females. Twenty surrogate variables were included as covariates to account for cell count heterogeneity and technical variation. These were calculated from the DNA methylation profiles using the sva R package (23).

Following the EWAS of age in each dataset, correlations between CpG site effect sizes revealed that six datasets had extremely low agreement with the other datasets (Supplementary Table 7; mean correlation < 0.25). We therefore excluded these datasets from meta-analysis. EWAS age associations were meta-analysed using fixed effect, inverse variance-weighted meta-analysis implemented by the metafor R package (24).

To apply dmrff in the meta-analysis setting, we first identified candidate regions using the meta-analysed EWAS p-values and effect sizes, i.e. regions defined by sets of CpG sites with consecutive CpG sites at most 500bp apart and each site having an EWAS p-value < 0.05 and the same EWAS effect sign. We then calculated the dmrff statistic for each region and sub-region in each meta-analysed dataset. Following this, the dmrff statistics from each dataset were then meta-analysed using fixed effect, inverse variance-weighted meta-analysis. Candidate DMRs were identified as in dmrff by greedily selecting sub-regions with the most extreme meta-analysed statistic that does not overlap with previously selected candidates. P-values for each candidate were Bonferroni-adjusted for multiple tests, conservatively treating each EWAS test and each sub-region test as independent.

This approach (called ‘two-step meta-analysis’), however, can be impractical when the meta-analysis is being performed by a consortium because it requires two interactions between the meta-analysis team and analysts of individual datasets. Dataset analysts first supply EWAS summary statistics and then must wait for the meta-analysis team to identify and communicate a set of genomic regions of interest so that they can calculate dmrff statistics for these regions for the final DMR meta-analysis. To reduce to a single interaction, we considered using a ‘reference’ dataset during meta-analysis to obtain CpG site correlations. To maximize agreement with the reference dataset, we selected the EWAS dataset that was most similar in terms of pairwise CpG site correlations to the other EWAS datasets. More specifically, we selected 10,000 random CpG sites and a random nearby mate and calculated the correlations between these pairs in each dataset. We then calculated the correlation of these correlations between all pairs of datasets. The reference dataset was selected as the one with the largest mean correlation with all other datasets. We found this to be dataset with GEO accession GSE72680 (mean R = 0.75; Supplementary Table 7). We call this approach ‘reference meta-analysis’.

## List of Abbreviations

CpG: Cytosine followed by a Guanine
DMR: Differentially Methylated Region
EPIC: Illumina Infinium MethylationEPIC BeadChip microarray
EWAS: Epigenome-Wide Association Study
FPR: False Positive Rate
GEO: Gene Expression Omnibus
450k: Illumina Infinium HumanMethylation450 BeadChip microarray

## Declarations

### Ethics approval and consent to participate

Not applicable.

### Consent for publication

Not applicable.

### Availability of data and material

All data is publicly available from the GEO website under the following accession numbers: GSE51057, GSE80417, GSE43414, GSE74548, GSE97362, GSE73103, GSE50660, GSE72775, GSE59065, GSE56046, GSE51032, GSE72773, GSE40279, GSE87571, GSE87648, GSE72680, GSE56105, GSE84727, GSE42861, GSE55763.

### Authors’ contributions

MS implemented the dmrff algorithm and oversaw data analyses. MS, JRS, AS, RF and KT designed the algorithm. MS, JRS, RF and RA performed analyses. All authors read, revised and approved the manuscript.

## Acknowledgements

Not applicable.

## Supplementary Materials

**Supplementary Table 1.** True positive rates (%) of each algorithm for different correlations.

| Correlations | 0 | 0.005 | 0.01 | 0.015 | 0.02 | 0.025 | 0.03 | 0.035 | 0.04 | 0.045 | 0.05 | 0.06 |
| --- | --- | --- | --- | --- | --- | --- | --- | --- | --- | --- | --- | --- |
| **bumphunter (area statistic)** | 0 | 0 | 0 | 0 | 0 | 0 | 0 | 0 | 0 | 0 | 0 | 0 |
| **bumphunter (value statistic)** | 0 | 0 | 0 | 0 | 0 | 0 | 0 | 0 | 0 | 0 | 0 | 0 |
| **comb-p** | 0 | 4 | 0 | 1 | 2 | 0 | 2 | 0 | 3 | 2 | 4 | 2 |
| **DMRcate** | 0 | 0 | 0 | 0 | 0 | 0 | 1 | 0 | 1 | 2 | 0 | 0 |
| **dmrff** | 0 | 0 | 0 | 0 | 0 | 0 | 1 | 0 | 0 | 1 | 0 | 0 |
| **EWAS** | 0 | 0 | 0 | 0 | 0 | 0 | 1 | 0 | 0 | 0 | 0 | 0 |
| **seqlm** | 0 | 0 | 0 | 0 | 0 | 0 | 0 | 0 | 0 | 0 | 0 | 0 |

| Correlations | 0.07 | 0.08 | 0.09 | 0.1 | 0.125 | 0.15 | 0.175 | 0.2 | 0.25 | 0.3 | 0.35 | 0.4 |
| --- | --- | --- | --- | --- | --- | --- | --- | --- | --- | --- | --- | --- |
| **bumphunter (area statistic)** | 0 | 0 | 0 | 0 | 0 | 0 | 0 | 0 | 1 | 3 | 4 | 3 |
| **bumphunter (value statistic)** | 0 | 0 | 0 | 0 | 0 | 0 | 0 | 2 | 5 | 10 | 13 | 12 |
| **comb-p** | 4 | 8 | 11 | 16 | 19 | 37 | 54 | 69 | 89 | 98 | 100 | 100 |
| **DMRcate** | 2 | 1 | 2 | 7 | 7 | 16 | 19 | 37 | 73 | 90 | 94 | 99 |
| **dmrff** | 0 | 0 | 0 | 1 | 2 | 7 | 13 | 21 | 72 | 90 | 97 | 100 |
| **EWAS** | 0 | 1 | 0 | 1 | 1 | 4 | 7 | 12 | 39 | 78 | 80 | 96 |
| **seqlm** | 0 | 1 | 0 | 2 | 4 | 7 | 14 | 34 | 71 | 96 | 98 | 100 |

**Supplementary Table 2.** Numbers of age DMRs identified in multiple datasets. Table ordered by the number of associations identified by EWAS.

| dataset | dataset<br>size | EWAS | EWAS<br>regions* | dmrff | seqlm |
| --- | --- | --- | --- | --- | --- |
| GSE50660 | 464 | 1208 | 1006 | 1235 | 978 |
| GSE59065 | 97 | 3014 | 2453 | 4156 | 0 |
| GSE72775 | 335 | 3243 | 2567 | 3291 | 1872 |
| GSE72773 | 310 | 4949 | 3869 | 4884 | 2459 |
| GSE51032 | 845 | 5310 | 3851 | 4892 | 4191 |
| GSE40279 | 656 | 10356 | 7345 | 9489 | 4736 |
| GSE56046 | 1202 | 10467 | 8143 | 9625 | 8334 |
| GSE87648 | 382 | 11348 | 7446 | 10488 | 6115 |
| GSE42861 | 689 | 14462 | 9024 | 12892 | 10320 |
| GSE84727 | 665 | 15132 | 9988 | 13243 | 9859 |
| GSE72680 | 422 | 18292 | 10533 | 16592 | 11623 |
| GSE56105 | 613 | 43010 | 25399 | 37434 | 16460 |
| GSE55763 | 2711 | 58906 | 30812 | 48975 | 46359 |
| GSE87571 | 729 | 112905 | 62068 | 97682 | 90931 |

\* EWAS regions are defined in terms of CpG sites associated with age (Bonferroni-adjusted p-value < 0.05) with at most 500bp between consecutive sites in the genome.

**Supplementary Table 3.** ‘Unique’ age DMRs identified in multiple datasets. The table is ordered by the sizes of the datasets. Each pair of algorithms is compared, and the DMRs identified by each algorithm (at Bonferroni-adjusted p < 0.05) that do not overlap with those identified by the other algorithm (at Bonferroni-adjusted p < 0.2) are shown.

| dataset | dataset size | dmrff vs EWAS |  |  | seqlm vs EWAS |  |  | dmrff vs seqlm |  |
| --- | --- | --- | --- | --- | --- | --- | --- | --- | --- |
|  |  | dmrff | EWAS regions* | EWAS | seqlm | EWAS regions* | EWAS | dmrff | seqlm |
| GSE59065 | 97 | 440 | 0 | 0 | 0 | 2453 | 3014 | 4156 | 0 |
| GSE72773 | 310 | 95 | 0 | 0 | 60 | 1252 | 1491 | 1590 | 28 |
| GSE72775 | 335 | 166 | 0 | 0 | 192 | 710 | 849 | 926 | 80 |
| GSE87648 | 382 | 622 | 0 | 0 | 112 | 1691 | 2231 | 2617 | 69 |
| GSE72680 | 422 | 115 | 0 | 0 | 182 | 1103 | 1685 | 1471 | 99 |
| GSE50660 | 464 | 64 | 0 | 0 | 69 | 127 | 154 | 195 | 34 |
| GSE56105 | 613 | 358 | 0 | 0 | 1 | 11487 | 16019 | 14700 | 0 |
| GSE40279 | 656 | 185 | 0 | 0 | 27 | 2663 | 3439 | 3266 | 5 |
| GSE84727 | 665 | 346 | 0 | 0 | 193 | 1576 | 2131 | 1813 | 43 |
| GSE42861 | 689 | 181 | 0 | 0 | 323 | 779 | 1118 | 1083 | 190 |
| GSE87571 | 729 | 379 | 0 | 0 | 582 | 1141 | 2631 | 2333 | 326 |
| GSE51032 | 845 | 133 | 0 | 0 | 269 | 372 | 494 | 502 | 146 |
| GSE56046 | 1202 | 249 | 0 | 0 | 144 | 507 | 702 | 764 | 23 |
| GSE55763 | 2711 | 605 | 0 | 0 | 445 | 1325 | 3088 | 2841 | 126 |

\* EWAS regions are defined in terms of CpG sites associated with age (Bonferroni-adjusted $p$ -value $< 0.05$ ) with at most 500bp between consecutive sites in the genome.

**Supplementary Table 4.**
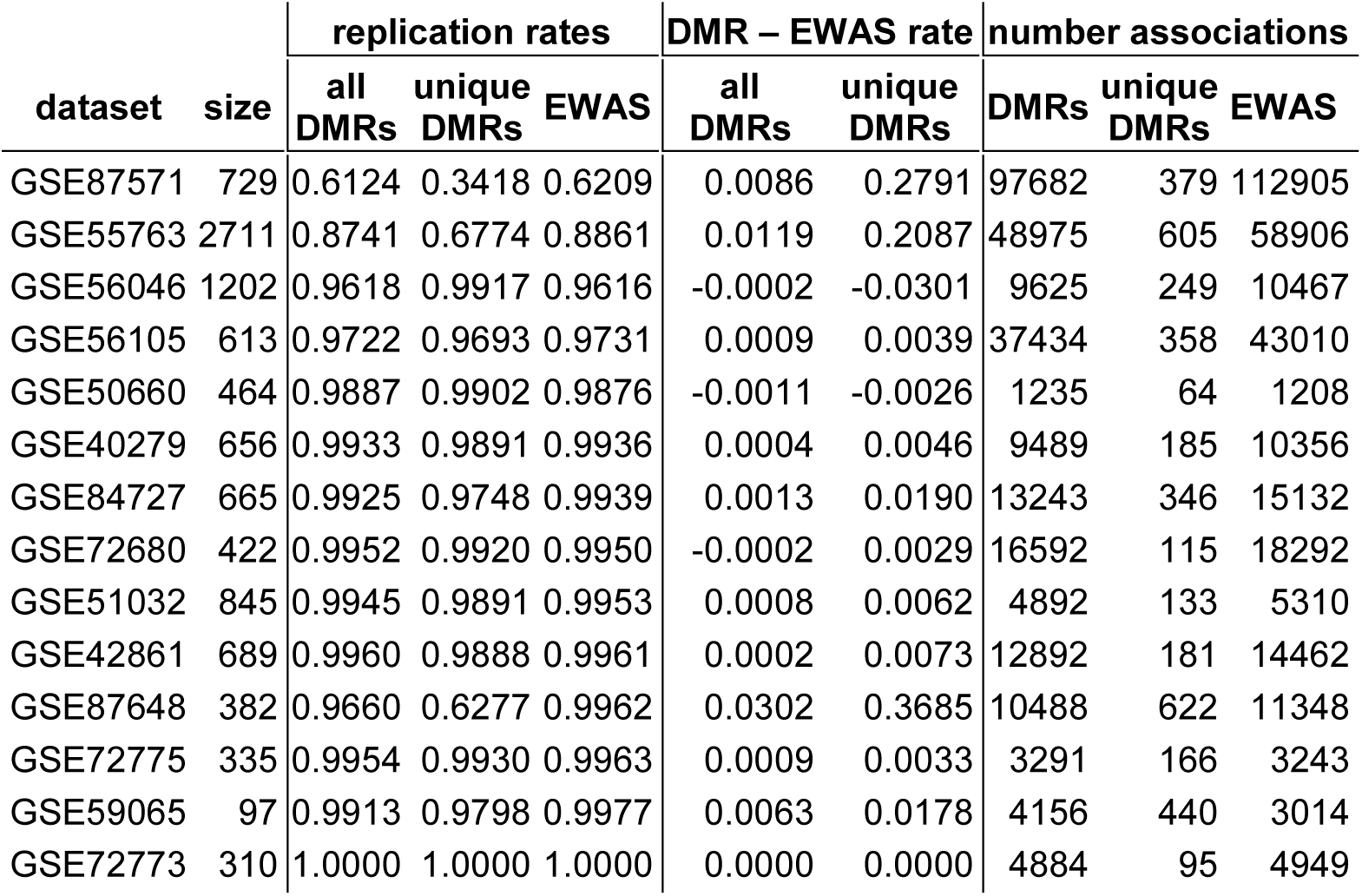
Replication rates of dmrff and EWAS in multiple datasets. Replication rates were calculated by meta-analysing the DMRs or associations observed in each dataset in the other datasets. ‘Unique’ DMRs are those that do not contain an EWAS association (at Bonferroni adjusted p < 0.2).

| dataset | size | replication rates |  |  | DMR – EWAS rate |  | number associations |  |  |
| --- | --- | --- | --- | --- | --- | --- | --- | --- | --- |
|  |  | all DMRs | unique DMRs | EWAS | all DMRs | unique DMRs | DMRs | unique DMRs | EWAS |
| GSE87571 | 729 | 0.6124 | 0.3418 | 0.6209 | 0.0086 | 0.2791 | 97682 | 379 | 112905 |
| GSE55763 | 2711 | 0.8741 | 0.6774 | 0.8861 | 0.0119 | 0.2087 | 48975 | 605 | 58906 |
| GSE56046 | 1202 | 0.9618 | 0.9917 | 0.9616 | -0.0002 | -0.0301 | 9625 | 249 | 10467 |
| GSE56105 | 613 | 0.9722 | 0.9693 | 0.9731 | 0.0009 | 0.0039 | 37434 | 358 | 43010 |
| GSE50660 | 464 | 0.9887 | 0.9902 | 0.9876 | -0.0011 | -0.0026 | 1235 | 64 | 1208 |
| GSE40279 | 656 | 0.9933 | 0.9891 | 0.9936 | 0.0004 | 0.0046 | 9489 | 185 | 10356 |
| GSE84727 | 665 | 0.9925 | 0.9748 | 0.9939 | 0.0013 | 0.0190 | 13243 | 346 | 15132 |
| GSE72680 | 422 | 0.9952 | 0.9920 | 0.9950 | -0.0002 | 0.0029 | 16592 | 115 | 18292 |
| GSE51032 | 845 | 0.9945 | 0.9891 | 0.9953 | 0.0008 | 0.0062 | 4892 | 133 | 5310 |
| GSE42861 | 689 | 0.9960 | 0.9888 | 0.9961 | 0.0002 | 0.0073 | 12892 | 181 | 14462 |
| GSE87648 | 382 | 0.9660 | 0.6277 | 0.9962 | 0.0302 | 0.3685 | 10488 | 622 | 11348 |
| GSE72775 | 335 | 0.9954 | 0.9930 | 0.9963 | 0.0009 | 0.0033 | 3291 | 166 | 3243 |
| GSE59065 | 97 | 0.9913 | 0.9798 | 0.9977 | 0.0063 | 0.0178 | 4156 | 440 | 3014 |
| GSE72773 | 310 | 1.0000 | 1.0000 | 1.0000 | 0.0000 | 0.0000 | 4884 | 95 | 4949 |

**Supplementary Table 5.**
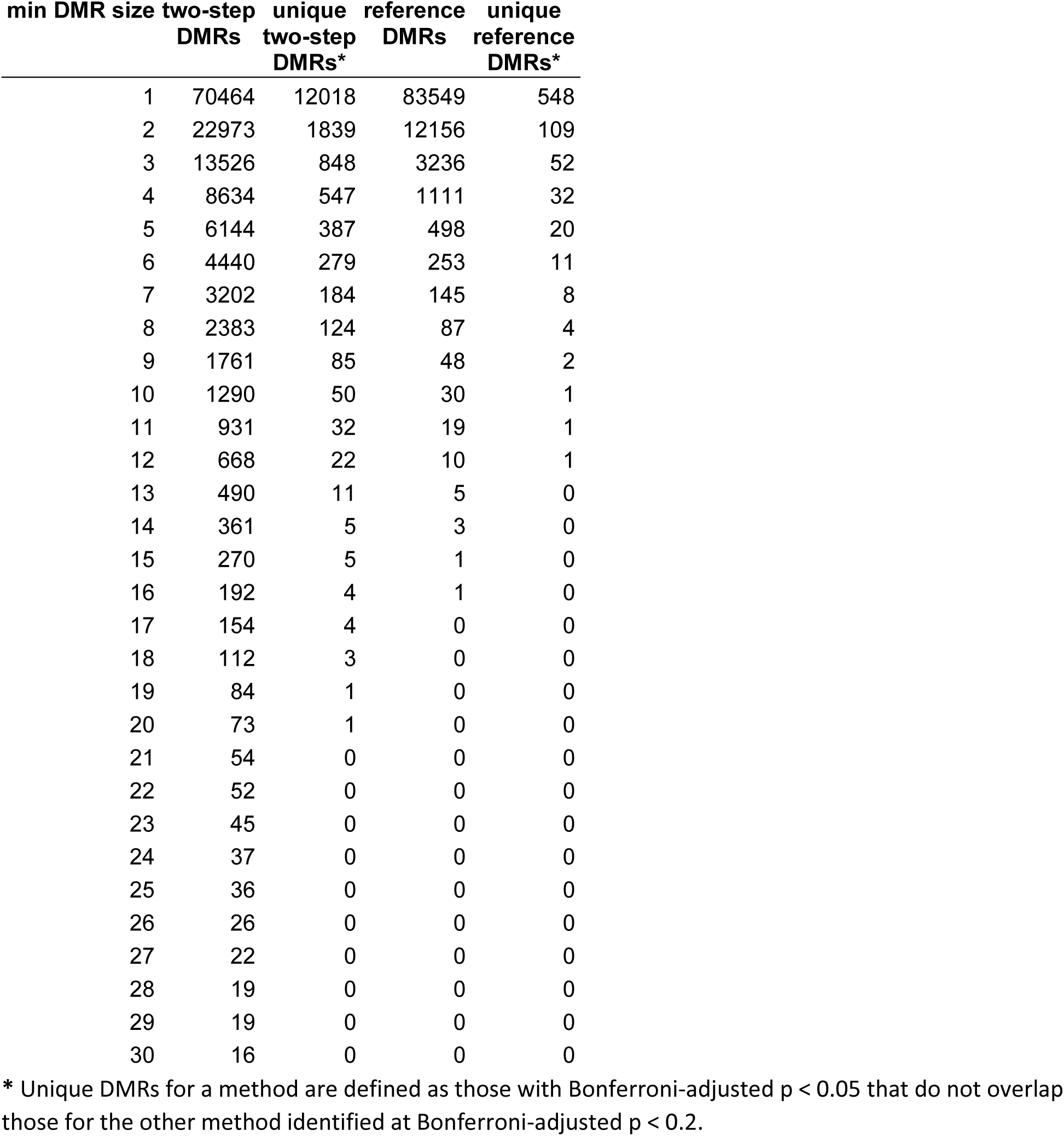
‘Unique’ age DMRs by numbers of CpG sites identified by the *full* two-step and reference dmrff meta-analysis methods.

| min DMR size | two-step DMRs | unique two-step DMRs* | reference DMRs | unique reference DMRs* |
| --- | --- | --- | --- | --- |
| 1 | 70464 | 12018 | 83549 | 548 |
| 2 | 22973 | 1839 | 12156 | 109 |
| 3 | 13526 | 848 | 3236 | 52 |
| 4 | 8634 | 547 | 1111 | 32 |
| 5 | 6144 | 387 | 498 | 20 |
| 6 | 4440 | 279 | 253 | 11 |
| 7 | 3202 | 184 | 145 | 8 |
| 8 | 2383 | 124 | 87 | 4 |
| 9 | 1761 | 85 | 48 | 2 |
| 10 | 1290 | 50 | 30 | 1 |
| 11 | 931 | 32 | 19 | 1 |
| 12 | 668 | 22 | 10 | 1 |
| 13 | 490 | 11 | 5 | 0 |
| 14 | 361 | 5 | 3 | 0 |
| 15 | 270 | 5 | 1 | 0 |
| 16 | 192 | 4 | 1 | 0 |
| 17 | 154 | 4 | 0 | 0 |
| 18 | 112 | 3 | 0 | 0 |
| 19 | 84 | 1 | 0 | 0 |
| 20 | 73 | 1 | 0 | 0 |
| 21 | 54 | 0 | 0 | 0 |
| 22 | 52 | 0 | 0 | 0 |
| 23 | 45 | 0 | 0 | 0 |
| 24 | 37 | 0 | 0 | 0 |
| 25 | 36 | 0 | 0 | 0 |
| 26 | 26 | 0 | 0 | 0 |
| 27 | 22 | 0 | 0 | 0 |
| 28 | 19 | 0 | 0 | 0 |
| 29 | 19 | 0 | 0 | 0 |
| 30 | 16 | 0 | 0 | 0 |

\* Unique DMRs for a method are defined as those with Bonferroni-adjusted $p < 0.05$ that do not overlap those for the other method identified at Bonferroni-adjusted $p < 0.2$ .

**Supplementary Table 6.** Publicly available datasets available for the age meta-analysis. For each dataset, the table provides the Gene Expression Omnibus accession, the number of samples, and the average correlation of EWAS age effects with those of the other datasets.

| **GEO accession** | **n** | **R** |
| --- | --- | --- |
| GSE51057 | 329 | 0.06 |
| GSE80417 | 638 | 0.07 |
| GSE43414 | 80 | 0.13 |
| GSE74548 | 174 | 0.15 |
| GSE97362 | 233 | 0.19 |
| GSE73103 | 355 | 0.24 |
| GSE50660 | 464 | 0.42 |
| GSE72775 | 335 | 0.46 |
| GSE59065 | 97 | 0.48 |
| GSE56046 | 1202 | 0.49 |
| GSE51032 | 845 | 0.51 |
| GSE72773 | 310 | 0.52 |
| GSE40279 | 656 | 0.53 |
| GSE87571 | 729 | 0.53 |
| GSE87648 | 382 | 0.54 |
| GSE72680 | 422 | 0.55 |
| GSE56105 | 613 | 0.55 |
| GSE84727 | 665 | 0.55 |
| GSE42861 | 689 | 0.56 |
| GSE55763 | 2711 | 0.58 |

**Supplementary Table 7.** Publicly available datasets used in the age meta-analysis. For each dataset, the table provides the Gene Expression Omnibus accession, the number of samples, and the average correlation of the estimated correlation structure within each dataset with that of the other datasets. Correlation structure was estimated within a dataset as the correlations between 10,000 randomly selected pairs of CpG sites.

| **GEO accession** | **n** | **R** |
| --- | --- | --- |
| GSE87648 | 382 | 0.53 |
| GSE56046 | 1202 | 0.61 |
| GSE50660 | 464 | 0.61 |
| GSE59065 | 97 | 0.63 |
| GSE72773 | 310 | 0.65 |
| GSE72775 | 335 | 0.67 |
| GSE42861 | 689 | 0.67 |
| GSE55763 | 2711 | 0.69 |
| GSE84727 | 665 | 0.70 |
| GSE51032 | 845 | 0.71 |
| GSE87571 | 729 | 0.72 |
| GSE56105 | 613 | 0.72 |
| GSE40279 | 656 | 0.72 |
| GSE72680 | 422 | 0.75 |

